## Supplementary Material for "Predictive learning as a network mechanism for extracting low-dimensional latent space representations"

#### Contents

**1 Predictive learning and representations in the simple "card game" example: Further analysis 1**
1.1 Learning neural representations in the card game example . . . . . . . . . . . . 1
1.2 Analysis of the regularity of representations . . . . . . . . . . . . . . . . . . . . . 2

**2 Theoretical analysis of predictive learning and latent space representations 3**
2.1 Low-dimensional neural representation manifolds and how they code latent variables 3
2.2 Linear Dimensionality analysis: Participation Ratio and Dimensionality Gain . . 5

#### 1 Predictive learning and representations in the simple "card game" example: Further analysis

##### 1.1 Learning neural representations in the card game example

In Fig. S1 we show how the neural representation, projected in the Principal Component space of PCs 1-2, develops throughout learning for the card game task. In Fig. S1a we show the learning progression for the same data as in plot of Fig.1d in the main manuscript. In Fig. S1b we color the same plot by the previous state while in Fig. S1c we color the plot by the action.

These plots show how the grid of states is a prospective grid. This means that the states represented in it are not the states of input of the network but rather states of output. This means that the latent structure extracted by the network is the latent structure of the outputs and not the inputs. These have the same latent structure in terms of lattice ordering but the points that are in proximity are not the ones that are generated with the same observation $o_s$ as input but rather with the same observation as output. This is a critical difference in the predictive learning representation, as we explore further below.

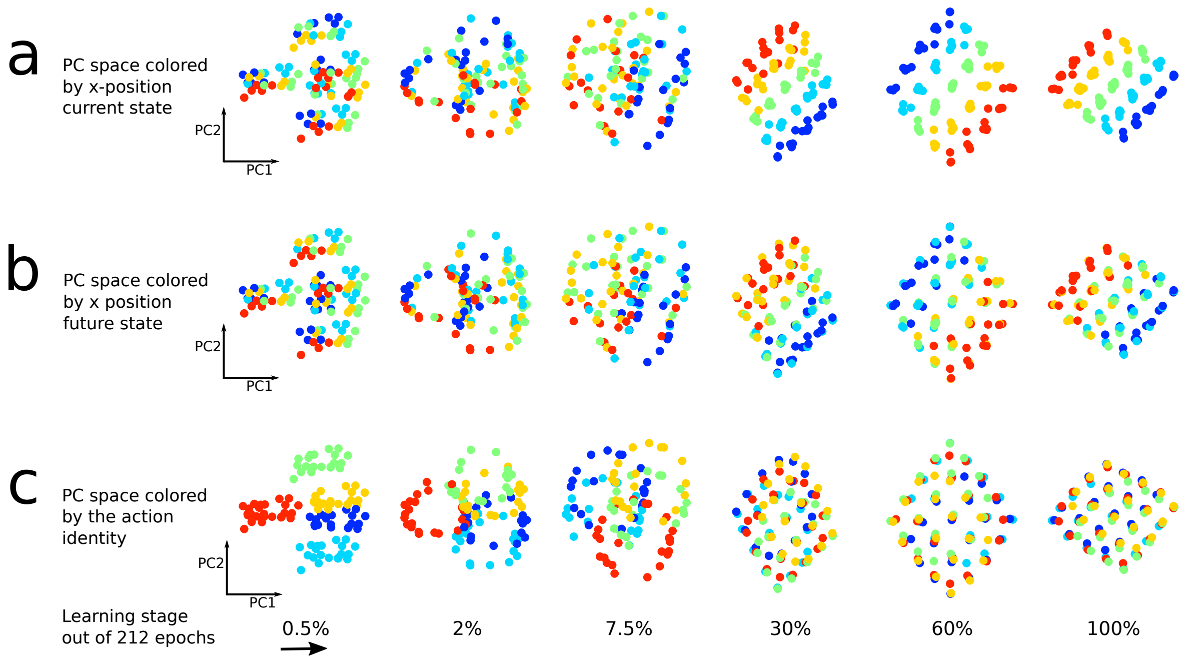

**Figure S1.** Learning the predictive neural representation a) Principal component space of the neural representation colored by the x-coordinate of the input latent space. b) Principal component space of the neural representation colored by the x-coordinate of the output latent space. c) Principal component space of the neural representation colored by the input action.

#### 1.2 Analysis of the regularity of representations

In Fig.1 of the main manuscript we show that, even when the underlying network is trained without actions, its representations still develop some regularity, but less than in the case when actions are provided. We here quantify this regularity. To do so we analyse Euclidean distances between the representations of different points in the network trained with and without actions. We compute this as a function of the state distance on the 2d lattice, where nearby states are considered to be at distance 1 while further states follow the Euclidean distance on the lattice. For example starting from a state and taking 2 move right (East) and one up (North) leads to a second state at a distance $\sqrt{5}$ from the original. In Fig. S2a we show the distributions of distances between the representations of all states at distance 2. The representation with actions displays a smaller variance and a higher average. In Fig. S2b we show the scaling of the average norms as a function of the distance between states. We see that the scaling in the network trained with actions appears perfectly linear. The fact that the scaling of distances in the network trained without actions also displays a linear relationship is indicative of the fact that the representation is "partially ordered." The quantification of this partial ordering or *regularity* is given in Fig. S2c where we show the average divided by the standard deviation of the distances between states (these are averages and standard deviations of all distance distributions as in Fig. S2a). We highlight how, in the network with actions, the linear trend is maintained, following from the fact that while the norm increases (Fig. S2b) the standard deviation is fairly constant. By contrast, for the network trained without actions, the standard deviation (the "noise" in this analysis) increases so that the relative increase in the average norm (the "signal") is damped.

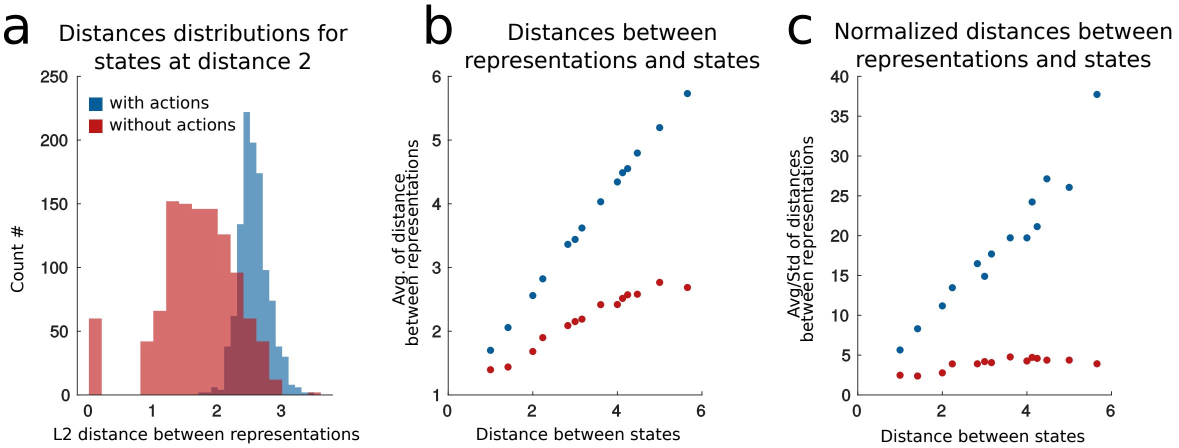

**Figure S2.** Regularity analysis. a) Distribution of distances for the representation of states at a lattice distance of 2 from one another. b) Average of the distribution of Euclidean distances of neural representations as a function of the distances between the two corresponding states. c) Same as b) but normalized by the standard deviation of each representation, i.e. displaying mean/std.

### 2 Theoretical analysis of predictive learning and latent space representations

#### 2.1 Low-dimensional neural representation manifolds and how they code latent variables

We begin by defining and characterising the dimensionality of a representation manifold in an idealized, pre-prescribed setting. This is a simplified, concrete model of latent space coding. Low-dimensional (Low-D) representation manifolds occur when a large number of neurons are strongly and consistently tuned to a small set of latent variables. Place and grid cells are examples of such coding [13, 18–20].

In the following, we consider the following specific setting. Given two continuous variables $x, y$ that parametrize a latent space, Fig. S3a, consider an ensemble of $N$ neurons with Gaussian tuning curves that are centered over uniformly distributed locations on the latent space. For example a neuron may be centered at location $(x_0, y_0)$ and have a gaussian radial basis tuning curve as shown in Fig. S3b, $\mathcal{G}_\sigma(x, y) = \frac{1}{2\pi\sigma^2} \exp\left(-\frac{(x-x_0)^2+(y-y_0)^2}{2\sigma^2}\right)$. The responses of an ensemble of $N$ neurons map the latent space manifold (Fig. S3a) to a neural response manifold embedded in neural representation space (that is, the $N$-dimensional space spanned by the activity of all neurons in the population. To visualize the response manifold, we project it onto its first three Principal Components (PCs), Fig. S3c. As the agent traverses a trajectory $\boldsymbol{x}_t$ in the 2d latent space (Fig. S3a, grayscale), the representation $\boldsymbol{r}_t$ traces out a trajectory on the response manifold (Fig. S3c, grayscale). We can view the tuning curve of a single neuron (Fig. S3b) on the response manifold to obtain the *manifold tuning curve* of this neuron (Fig. S3d), as in Fig. 5 in the main text. In the next section we will analyze in more depth the meaning and properties of manifold tuning curves.

The two dimensions of the latent space completely parametrize the response manifold, resulting in a two-dimensional curved surface. The fact that the representation manifold has two dimensions is revealed by a measure known as Intrinsic Dimensionality (ID), whose formal definition relies on concepts of Riemannian geometry for smooth manifolds [5].

While the ID of the representation manifold is two, due to its curvature, many linear components are necessary to cover it in the $N$-dimensional neural space. This linear dimensionality can be captured by a second measure of dimensionality: the Participation Ratio (PR) of the

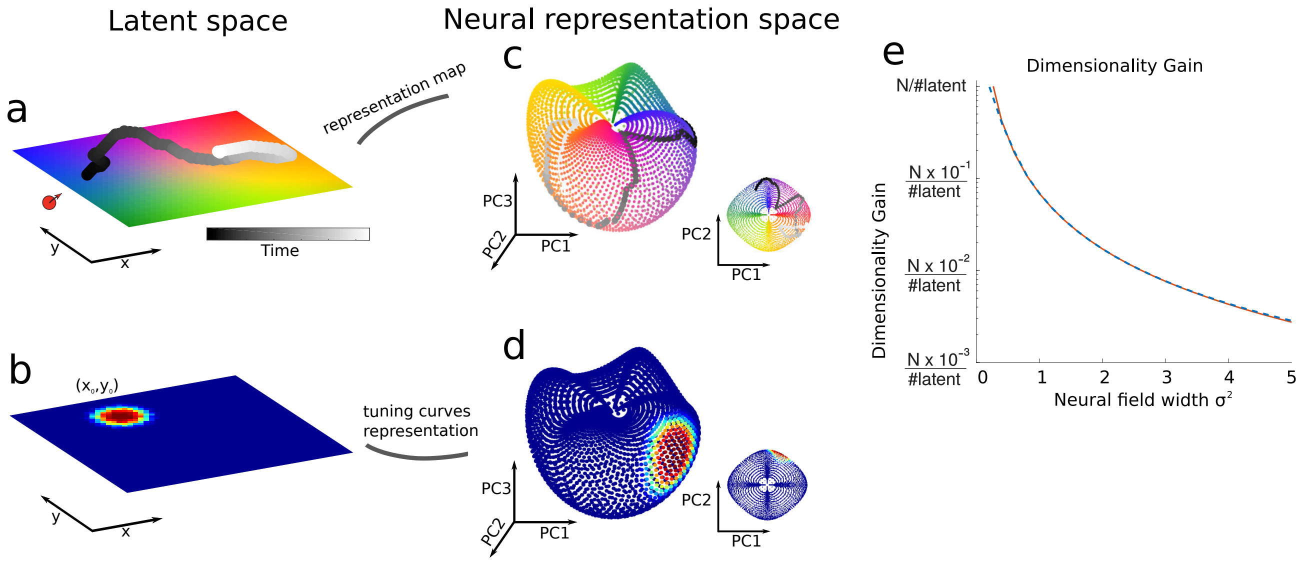

**Figure S3.** Analysis of neural representation manifolds with pre-prescribed neural tuning. a) Example of a two dimensional environment in which the agent moves. We assign a unique color to each location of the environment. A segment of the agent's trajectory is represented in gray scale, with shade standing for time. b) Example tuning of a neuron with gaussian receptive field centered on $(x_0, y_0)$. c) Neural representation manifold projected onto PCs 1 to 3, under the assumptions that neurons have gaussian receptive fields which uniformly cover the environment and that the agent uniformly explores the environment. Displayed points are uniformly sampled from the manifold. Each point of this representation manifold is colored according to the corresponding location in latent space. The agent's trajectory is represented on the manifold; the inset shows the top view (first two PCs). d) Example of a neural response field on the manifold. The same neuron shown in b) is now shown, with its receptive field with respect to manifold coordinates. e) Dimensionality Gain dependence on the size of the gaussian field $\sigma$. The red line represents the DG as computed for 4096 neurons tiling the latent space. The blue dotted line represents the theoretical analysis. In this case DG = PR/2 as the Intrisic Dimensionality ID = 2.

manifold. This metric is defined over the eigenvalues $\lambda_{1..N}$ of the covariance matrix $\boldsymbol{C}$ of the neural activity:

$$\mathrm{PR} = \frac{(Tr\boldsymbol{C})^2}{Tr(\boldsymbol{C}^2)} = \frac{(\sum_{i=1}^{N}\lambda_i)^2}{\sum_{i=1}^{N}\lambda_i^2} = \frac{1}{\sum_{i=1}^{N}\tilde{\lambda}_i^2} \tag{1}$$

where $\tilde{\lambda}_i = \lambda_i / \sum_{j=1}^{N}\lambda_j$, see Fig. S4a. [1, 7, 9, 12].

The two most important aspects of these measures of dimension are:

- ID of the representation manifold is determined by the latent variables underlying the inputs. As such, it does not depend on specific details of the neural code.
- PR, by contrast, is a property of the neural code. The more *localized* the neural fields are (i.e. the smaller the response curve width $\sigma$ is), the more decorrelated the neural activations are, and, in turn, the higher the linear dimensionality PR is.

Thus, the difference between PR and ID carries information about the non-linear embedding of latent variables in the representation. We suggest a novel metric, *Dimensionality Gain* (DG), to capture such difference which measures the extent to which a given representation linearly expands the "true" (i.e. intrinsic) dimensionality of the manifold:

$$\mathrm{DG} = \frac{\textit{linear dimensionality measure}}{\textit{non-linear dimensionality measure}} = \frac{\mathrm{PR}}{\mathrm{ID}} \,. \tag{2}$$

Fig. S3e shows a key observation, that we will return to in the context of predictive representations: that the Dimensionality Gain (DG) increases as the width $\sigma$ of the neural fields decreases. Thus a higher DG is regarded as a signature of low-D coding. We now give an analytical formula for this relationship as well as a more thorough explanation of relationships among ID, PR, and DG.

#### 2.2 Linear Dimensionality analysis: Participation Ratio and Dimensionality Gain

Participation Ratio is a measure of dimensionality that is based on the distributions of eigenvalues $(\lambda_1, \lambda_2 ...)$ of the covariance matrix $\boldsymbol{C}$:

$$PR = \frac{(\mathrm{Tr}\boldsymbol{C})^2}{\mathrm{Tr}(\boldsymbol{C}^2)} = \frac{(\sum_{i=1}^{N} \lambda_i)^2}{\sum_{i=1}^{N} \lambda_i^2} = \frac{1}{\sum_{i=1}^{N} \tilde{\lambda}_i^2} \tag{3}$$

where $\tilde{\lambda}_i = \lambda_i / \sum_{j=1}^{N} \lambda_j$. In the case of the example of Fig. S3, if we assume that all the locations of the latent space $\mathcal{X}$ are visited with the same probability, then we can compute the covariance matrix of the representation $\boldsymbol{C}$. The entry of the covariance matrix that corresponds to two neurons, $i$ and $j$, with neural fields centered respectively in position $\boldsymbol{x}_i \equiv (x_i, y_i)$ and $\boldsymbol{x}_j \equiv (x_j, y_j) = \boldsymbol{x}_j + \Delta\boldsymbol{x} = (x_i + \Delta x, y_i + \Delta y)$ and with isotropic variance $\boldsymbol{\sigma}^2 \equiv (\sigma_x^2, \sigma_y^2) = (\sigma^2, \sigma^2)$ is given by:

$$\boldsymbol{C}_{ij} = \frac{1}{T}\int_0^T dt\ (\mathcal{G}_\sigma(\boldsymbol{x}_i - \boldsymbol{x}_t) - \frac{1}{T}\int_0^T \mathcal{G}_\sigma(\boldsymbol{x}_i - \boldsymbol{x}_s)ds)(\mathcal{G}_\sigma(\boldsymbol{x}_j - \boldsymbol{x}_t) - \frac{1}{T}\int_0^T \mathcal{G}_\sigma(\boldsymbol{x}_j - \boldsymbol{x}_s)ds) \tag{4}$$

As each location of the latent space is visited uniformly then this time integral is equivalent to a spatial average over the area $A$ of the latent space $\mathcal{X}$:

$$\begin{aligned}\boldsymbol{C}_{ij} &= \frac{1}{A}\int_A dt(\mathcal{G}_\sigma(\boldsymbol{x}_i - \boldsymbol{x}_t) - \frac{1}{A})(\mathcal{G}_\sigma(\boldsymbol{x}_j - \boldsymbol{x}_t) - \frac{1}{A}) = \frac{1}{A}\int_A dt\ \mathcal{G}_\sigma(\boldsymbol{x}_i - \boldsymbol{x}_t)\mathcal{G}_\sigma(\boldsymbol{x}_j - \boldsymbol{x}_t) - \frac{1}{A} = \\ &= \frac{1}{4\pi\sigma^2}\frac{1}{A}e^{-\frac{\Delta^2}{4\sigma^2}}\int_A dt\ \mathcal{G}_{\sigma/\sqrt{2}}((\boldsymbol{x}_i + \boldsymbol{x}_j)/2 - \boldsymbol{x}_t) - \frac{1}{A} = \\ &= \frac{1}{4\pi\sigma^2 A}e^{-\frac{\Delta^2}{4\sigma^2}} - \frac{1}{A^2}\ .\end{aligned} \tag{5}$$

where we recall that $\mathcal{G}_\sigma$ is a Gaussian with variance $\sigma^2$ normalized to 1 over the area $A$. Eq. 5 shows that $C_{ij}$ has a banded structure; in particular it is in Toeplitz form, with entries that decay with the distance between neurons in latent space [7].

We can now compute the terms in Eq. 3 that determine the PR. Specifically by considering the approximation $A \gg 4\pi\sigma^2$ we obtain:

$$\begin{aligned}(\boldsymbol{C}^2)_{ij} &= \sum_{k=1}^{N} C_{ik}C_{jk} \approx \int_A \mathcal{G}_\sigma(i-k)\mathcal{G}_\sigma(k-j)dk = \\ &= \frac{1}{8\pi^2\sigma^2 A}e^{-\frac{\Delta_{ij}^2}{8\sigma^2}}\ .\end{aligned} \tag{6}$$

Thus the PR in the limit of large $N$ is:

$$PR = \frac{(Tr\boldsymbol{C})^2}{Tr(\boldsymbol{C}^2)} = \left(\frac{N}{4\pi\sigma^2 A}\right)^2 \frac{8\pi^2\sigma^2 A}{N} = \frac{NA}{2\pi\sigma^2}\ . \tag{7}$$

This shows that the PR dimensionality grows with the inverse of the width of the Gaussian kernel and is proportional to the number of neurons N. Furthermore we also see that it scales as

$\frac{A}{s\pi\sigma^2}$ which is the area divided by the width of the field which matches the intuition of the problem.

If all the principal components of neural representations are independent and have equal variance, all the eigenvalues of the covariance matrix have the same value and PR($\boldsymbol{C}$) $= N$. Alternatively, if the components are correlated so that the variance is evenly spread across M dimensions, then $\lambda_1 = \lambda_2 = \lambda_3 = ...\lambda_M$ with $\lambda_M > 0$ and $\lambda_m = 0$ for $m > M$ so that the data points are arranged in an M-dimensional subspace of the full N-dimensional space. In this case only M eigenvalues would be nonzero and PR($\boldsymbol{C}$) $= M$ (Fig. S4a). For other cases, this measure interpolates between these two regimes. As a rule of thumb, [7] establishes that the PR dimensionality can be thought as the number of dimensions required to explain about 80% of the total population variance in many applications.

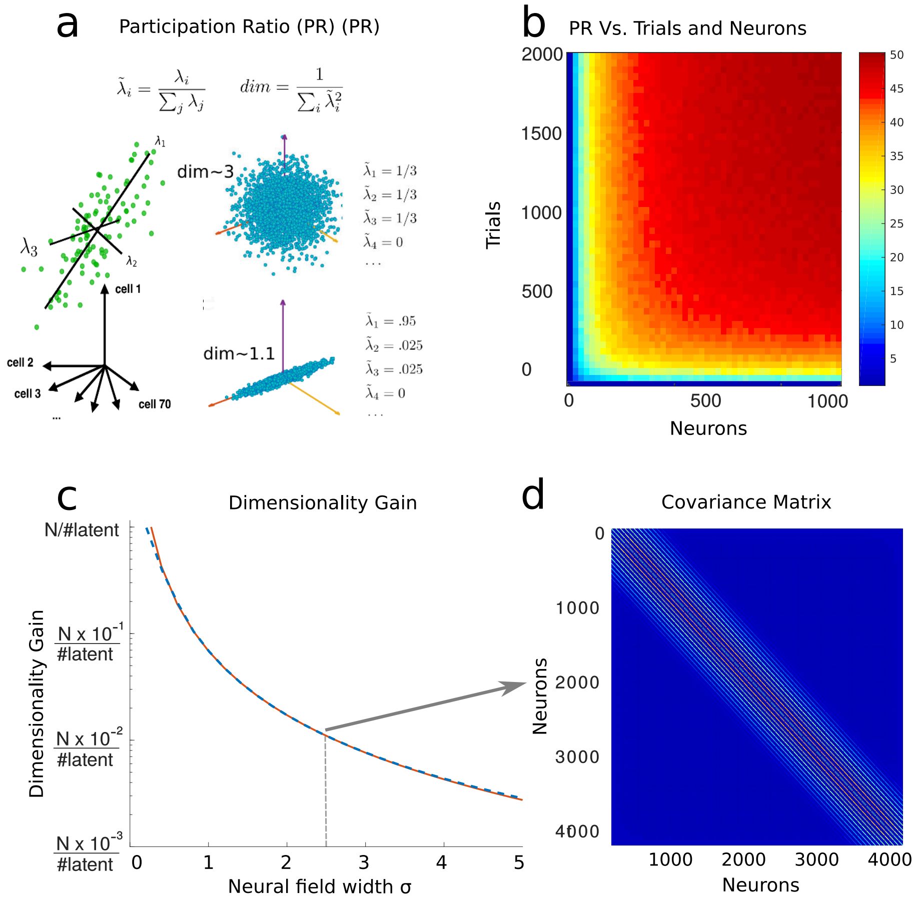

**Figure S4.** Linear dimensionality analysis. a) Illustration of the Participation Ratio (PR) dimensionality measure. The mathematical expression in terms of the eigenvalues of the covariance is illustrated for a few distributions in PC space. The left part shows an example of point cloud distribution and the leading eigenvalues $\lambda_{1,2,3}$. The right part shows a symmetric spherical distribution with PR=3 and an elongated one with PR=1.1. The eigenvalues of the covariance matrix are shown next to each example. b) PR estimation from a finite number of neurons or trials for the manifold example of Fig. S3 with $\sigma = 2.5$. c) PR dependence on the size of the gaussian field $\sigma^2$, same as figure Fig. S3e. The red line represents the DG as computed for 4096 neurons tiling the latent space shown in Fig.2 Main Text. The blue dotted line represents the theoretical analysis. d) Example of the covariance matrix for $\sigma = 2.5$.

#### 2.3 How latent space signal transfer follows from translation-invariant representations of neural states

This section explains the theory behind the results on latent space signal transfer shown in Figs. 3-6 of the main manuscript.

The analysis of the covariance matrix $\boldsymbol{C}$ developed above shows that it is in the Toeplitz form, due to the evenly spaced Gaussian tuning curves (cf. Fig. S4). Specifically, in the case shown in Fig. S4d, it is a Toeplitz tensor because, for each of the two variables, the Toeplitz structure is encoded in the representation as described in Fig. S3. Signal transfer measures the colinearity of the projection of the neural activity on the top eigenvectors of the covariance matrix with the latent variables. In the case analyzed above, $x$ and $y$ are the latent variables and signal transfer measures whether these two variables can be expressed as a linear combination of the projections on the top eigenvectors of the covariance matrix. To see when this is the case we need to compute the eigenvectors of the covariance in terms of $x$ and $y$. We first restrict our analysis to a nearly Toeplitz matrix in a single variable, Fig. S5a. The eigenvectors of such a Toeplitz matrix have recently been determined to be in a form approaching $\xi_i = a\ cos\left(\frac{\pi k i}{N+1}\right)$ for N large enough, where $a$ is the normalization coefficient and $k$ indicates the k$th$ eigenvector [3,4], Fig. S5b. The eigenvalues are shown in Fig. S5c which displays the relative importance of the first few eigenvectors, Fig. S5d.

The projections on the top eigenvectors are the elements of the representation that most contribute to the value of the signal transfer measure. The top eigenvector is the constant vector $\boldsymbol{n} = \frac{1}{\sqrt{N}}(1, 1, ..1)$. Projecting on this vector is equivalent to taking the average of the representation vector. In neuroscientific terms this would be the average activity or average firing rate across all neurons. The contribution of this eigenvector is subtracted when we consider a mean substracted covariance, the case displayed in the figure is for a Toeplitz matrix with rows normalized to have sum one rather than zero.

The second eigenvector follows the cosine function. Suppose as above that the response of the network to the inputs is similar to the response of a set of Gaussian-bump units responding selectively to the position latent variable $x$. Projecting the activity of the network onto the second eigenvector approximately returns the position at which the active bump is centered, but shifted by a constant and possibly negated (depending on the sign of the eigenvector). The reason for this is that projecting onto $\boldsymbol{\xi}$ such that $\xi_i = a\ cos\left(\frac{\pi i}{N+1}\right)$ for $i \in [1, N]$, is similar to projecting onto $\xi_i = -\frac{\pi i}{N+1} + 1$, since $\cos(x) \approx -x + 1$ in the interval $[0, \pi]$. Dropping the shift by $+1$, the magnitude of the correlation coefficient between $\cos(x)$ and $x$ in the interval $[0, \pi]$ is also large, equaling $\frac{4\sqrt{6}}{\pi^2} = 0.9927$, Fig. S5d (this is because $\cos(x)$ and $x$ are strongly anti-correlated).

Thus, if we assume a Gaussian response in the activity $f(\boldsymbol{x})$ with the form $f(\boldsymbol{x}) = \mathcal{G}_\sigma(\boldsymbol{x} - \boldsymbol{x}_0)$ around the true location $\boldsymbol{x}_0$ in the latent space $\mathcal{X}$, then the projection over the top eigenvectors of the covariance matrix in Toeplitz form returns a value strongly correlated (in magnitude) with the position $x_0$ of such gaussian bump that is the latent variable encoded by it. To see this consider the convolution, similar to this projection operation, between a gaussian and a linear variable:

$$\frac{1}{\sqrt{2\pi\sigma}} \int_{-\infty}^{+\infty} \mathcal{G}_\sigma(x - y)x\ dx = \frac{1}{\sqrt{2\pi\sigma}} \int_{-\infty}^{+\infty} e^{-\frac{(x-y)^2}{2\sigma^2}} \left((x - y) + y\right)\ dx = y\ . \tag{8}$$

This suggests that projecting over the PCs for a low-D code will lead to recovery of the latent variables. A condition for this to occur is that many cells are tuned to the underlying latent variables.

Now we consider the full case of the network responding to two position variables $x$ and $y$. The tensoring of multiple variables doesn't affect the argument above as the tensored space will have, as leading eigenvectors, the leading tensored eigenvectors of the individual spaces. The tensored covariance will be in the form:

$$\boldsymbol{C}_{xy} = \boldsymbol{C}_x \otimes \boldsymbol{C}_y$$

where the Kronecker tensor product is denoted by $\otimes$. Thus, for the case of two variables analyzed in depth in the previous section (Fig. S4), projecting on the first few eigenvectors still serves the role of recovering latent variables. For a deeper analysis and understanding of these phenomena we point the interested reader to more exhaustive reviews [3, 6, 16]. The most important caveat to this analysis is that the spectral properties of the Toeplitz matrix described above depend on the boundary conditions. The case we considered here, where the rows are normalized to sum to one, falls outside the common definition of Toeplitz matrix where the rows are truncated at the boundaries. This latter choice, with different boundary conditions, would lead to eigenvectors of the form $\xi_i = a\ sin\left(\frac{\pi k i}{N+1}\right)$ rather than $\xi_i = a\ cos\left(\frac{\pi k i}{N+1}\right)$, where $a$ is the normalization coefficient and $k$ indicates the k$th$ eigenvector. Thus, in this case the leading eigenvectors would be sine rather than cosine functions. This difference, however, doesn't interfere with the argument we illustrated above, although in this case is necessary to project on multiple eigevectors to reconstruct the latent variable. To this end a Canonical Correlation Analysis between the latent variables and the leading iegenvectors, as we perform in the main text in defining Latent Space Signal Transfer, comes in handy. For example, considring the canonical correlation coefficient between the underlying variable $x$ and the top four eigenvectors as sine functions ($k \in \{1, 2, 3, 4\}$) leads to a correlation coefficient of 0.86.

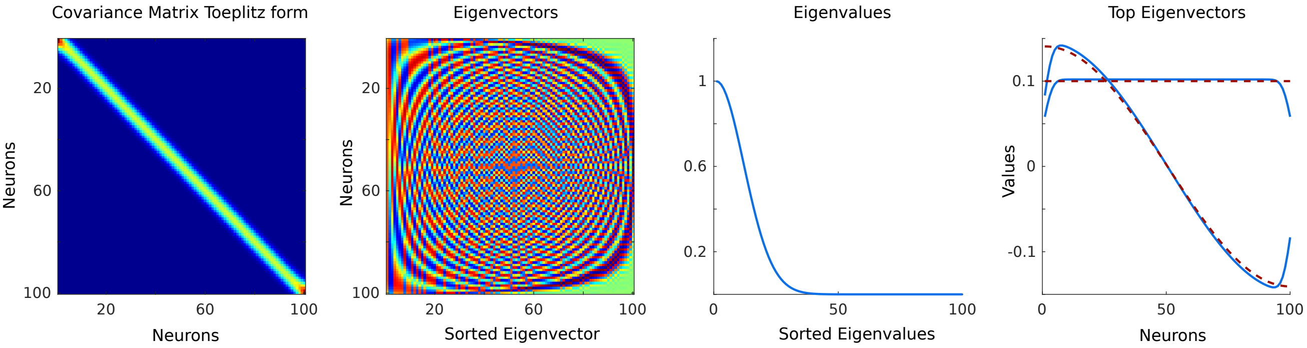

**Figure S5.** a) Covariance matrix in Toeplitz form. The normalization of the rows (summing up to one) is such that the boundary conditions for this matrix are not exactly in the Toeplitz. b) Sorted eigenvectors of the Toeplitz matrix in a). c) Sorted eigenvalues. d) Top two eigenvectors of the matrix: constant and cosine shaped. Numerical solutions are in blue and theory in red.

#### 2.4 Participation ratio and linear dimensionality

The arguments above imply that predictive representations will have low ID (i.e., low nonlinear dimensionality). We next give reasoning for why such predictive representations develop localized receptive fields. As shown in Fig. S3f, this leads, in turn, to high PR (i.e., high linear dimensionality) and hence high DG, all phenomena that we have observed in our network simulations above.

We begin with the assumption that the low-dimensional predictive representations are a smooth map of the latent space. A consequence is Lipschitz continuity, which guarantees that nearby points in the latent space $(\boldsymbol{x}, \boldsymbol{x}')$ map onto nearby points $(\boldsymbol{r}, \boldsymbol{r'})$ in representation space, at least up to a given radius:

$$d_{\boldsymbol{r},\boldsymbol{r'}} \leq \kappa d_{\boldsymbol{x},\boldsymbol{x'}} \tag{9}$$

where $\kappa$ is the Lipschitz constant and $d$ indicates distance. This preservation of distances, or similarities – together with the positivity constraint ($r_i \geq 0$ for each neuron i) – is known to lead to localized manifold fields [15, 17]. Interestingly, in our framework this result appears to be true for both positive representations (when the activation function is a sigmoid) and other ones although in such cases the localization of the receptive fields appears to be different and, in general, less localized than in the case where a sigmoid (positive) transfer function is used, Fig. S6.

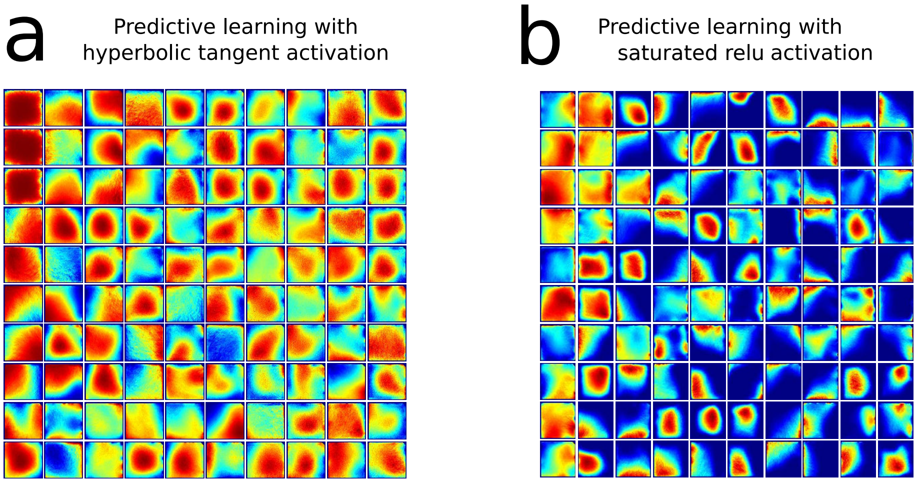

**Figure S6.** Neural activations comparison across activation functions. a) Average neural activations for a predictive network trained with hyperbolic tangent activation function. b) Same as panel a for a network trained with a hard sigmoid:
$f(x) = 0$ if x≤-2.5, $f(x) = 1$ if x≥2.5, $f(x) = 0.2x + 0.5$ otherwise.

The arguments above indicate that predictive learning leads to increases in linear dimensionality, as observed in our learning simulations (Fig.3 main manuscript). But when should this increase stop? A possible answer is: when the linear dimensionality of the neural representation matches that of the outputs that the network is seeking to produce. We give a simplified argument based on linear readout that suggests why this answer might be correct. Rewriting the cost function for a linear readout we obtain $\mathcal{C}_{pred} = \frac{1}{T}\sum_{t=0}^{T-1} ||\boldsymbol{o}_{t+1} - \boldsymbol{y}_t||^2 = \frac{1}{T}\sum_{t=0}^{T-1} ||\boldsymbol{o}_{t+1} - \boldsymbol{W}_{out}\boldsymbol{r}_t||^2$, and recognize that (for $\boldsymbol{W}_{out}$ randomly distributed or orthogonal), the linear dimensionality of the representation tends to match the linear dimensionality of the output as they are directly related through the linear transformation $\boldsymbol{W}_{out}$ (cf. [2,8,11]). Our numerical studies lend evidence to this: the PR increases through learning until it saturates at about the PR dimensionality of the output, which is 16.2, Fig.3 main manuscript.

#### 2.5 Further considerations on the locality of receptive fields

Consider the case where the movement of the agent in the latent space $\mathcal{X}$ is governed by a discrete-time dynamical system, similar to the case in the main text:

$$\boldsymbol{x}_{t+1} = F(\boldsymbol{x}_t) \tag{10}$$

where $\boldsymbol{x} = (x, y, \theta)$ and $F(\boldsymbol{x})$ is a vector field on $\mathcal{X}$. Above we argued that the recurrent network representation $\boldsymbol{r}_t = f^{RNN}(\boldsymbol{o_t}, \boldsymbol{r}_{t-1})$ through learning becomes a direct function of the latent space $\mathcal{X}$ as predictive learning extracts the latent variables: $\boldsymbol{r}_t = f(\boldsymbol{x_t})$. We now ask the question of whether this representation has localized neural activity.
Considering the local expansion at second order around a point $\boldsymbol{x}^* \in \mathcal{X}$ we obtain:

$$f(\boldsymbol{x}^*) - f(\boldsymbol{x}) = f(\boldsymbol{x}^*) + D_f(\boldsymbol{x}^*)\cdot(\boldsymbol{x} - \boldsymbol{x}^*) + (\boldsymbol{x} - \boldsymbol{x}^*)\cdot H_f(\boldsymbol{x}^*)\cdot(\boldsymbol{x} - \boldsymbol{x}^*) + ... \tag{11}$$

where $D_f$ and $H_f$ are respectively the Jacobian and Hessian. Assuming that the function $f$ is Lipschitz continous then:

$$d_{\boldsymbol{r}^*,\boldsymbol{r}} = ||f(\boldsymbol{x}^*) - f(\boldsymbol{x})|| \leq \kappa_m ||\boldsymbol{x} - \boldsymbol{x}^*|| \ , \tag{12}$$

where $\kappa_m$ is the Lipschitz constant. Furthermore if the inverse is Lipschitz, as expected if the representation manifold is smooth, then we have the bi-Lipschitz property:

$$\kappa_l d_{\boldsymbol{x},\boldsymbol{x}^*} \leq d_{\boldsymbol{r}^*,\boldsymbol{r}} = ||f(\boldsymbol{x}^*) - f(\boldsymbol{x})|| \leq \kappa_m d_{\boldsymbol{x},\boldsymbol{x}^*} \ . \tag{13}$$

These bounds suggest that local similarities in latent space $\mathcal{X}$ translate in local similarities in representation space $\mathcal{R}$. Furthermore, depending on the order of the Taylor series which dominates the local expansion of the function $f(\boldsymbol{x})$, we obtain a stronger form of Lipschitz continuity – Holder continuity:

$$\kappa_l d^{\alpha_l}_{\boldsymbol{x},\boldsymbol{x}^*} \leq d_{\boldsymbol{r}^*,\boldsymbol{r}} = ||f(\boldsymbol{x}^*) - f(\boldsymbol{x})|| \leq \kappa_m d^{\alpha_m}_{\boldsymbol{x},\boldsymbol{x}^*} \ . \tag{14}$$

These relationships control how representations of similar latent variables map onto similarities in the representation space, up to a certain radius. As latent variables become more and more distant, the corresponding representations tend to orthogonalize:

$$d^2_{\boldsymbol{x},\boldsymbol{x}^*} = ||\boldsymbol{x} - \boldsymbol{x}^*||^2 = ||\boldsymbol{x}||^2 + ||\boldsymbol{x}||^2 - 2\langle \boldsymbol{x}, \boldsymbol{x}^* \rangle \ , \tag{15}$$

which shows that as the scalar product $\langle \boldsymbol{x}, \boldsymbol{x}^* \rangle$ increases, the distance $d_{\boldsymbol{x},\boldsymbol{x}^*}$ decreases. On a spherical surface, where the norm $||\boldsymbol{x}||$ of each point is equal, the scalar product is in 1-1 correspondence with the distance.
An example of a code which varies continuously locally but orthogonalizes globally is a representation with localized gaussian fields, cf. Fig.2a-d in the main text. This phenomenon has been studied, with the extra condition of the representation being positive ($\boldsymbol{r} \geq 0$) in [17] where the authors show that preserving local similarities with a positivity constraint builds a representation whose receptive fields tile the representation manifold.
In sum, the arguments above indicate why activity on the representation manifold becomes localized in terms of the latent variables $\mathbf{x}$.

We close by emphasizing that the representations produced by the underlying neural networks will also be local in time. For example, consider a Wiener process in the latent space. If $\boldsymbol{x}_{t+1} = \boldsymbol{x}_t + \boldsymbol{\xi}$ and $\boldsymbol{\xi}$ is isotropically i.i.d. according to a Gaussian distribution $\mathcal{G}(0, \sigma^{\mathcal{X}})$ for each coordinate, then we obtain the relations:

$$d_{\boldsymbol{x(t)},\boldsymbol{x(t^*)}} = ||\boldsymbol{x}(t^*) - \boldsymbol{x}(t)|| = d^{\mathcal{X}} \sigma^{\mathcal{X}} \sqrt{t^* - t} \ , \tag{16}$$

where $d^{\mathcal{X}}$ is the dimensionality of the latent space. Such relations lead to

$$\kappa_l d^{\alpha_l}_{\boldsymbol{x},\boldsymbol{x}^*} = \kappa_l (d^{\mathcal{X}} \sigma^{\mathcal{X}} \sqrt{t^* - t})^{\alpha_l} \leq d_{\boldsymbol{r}^*,\boldsymbol{r}} = ||f(\boldsymbol{x}(t^*)) - f(\boldsymbol{x}(t))|| \leq \kappa_m d^{\alpha_m}_{\boldsymbol{x},\boldsymbol{x}^*} = \kappa_m (d^{\mathcal{X}} \sigma^{\mathcal{X}} \sqrt{t^* - t})^{\alpha_m} \ . \tag{17}$$

This equation highlights how similarities scale with time. They also scale with the dimensionality of the representation manifold $d^{\mathcal{R}}$, so that considering the effective random dynamics induced on it, we have:

$$d_{\boldsymbol{r_t},\boldsymbol{r_{t^*}}} \geq d^{\mathcal{R}} \sigma^{\mathcal{R}} \sqrt{t^* - t} \ . \tag{18}$$

Here $\sigma^{\mathcal{R}}$ denotes the average variance, per dimension, of the induced Wiener process in representation space. As the dimensionality of the manifold $d^{\mathcal{R}}$ decreases then the bounds become tighter and the similarity between neighbouring points increases. These considerations will drive future research aimed at fully describing how similarities explored dynamically across time lead to the learning of similarities across space on the representation manifold.

### 3 Control studies: Numerical simulations

#### 3.1 Robustness of our findings: comparing results for multiple tasks and network structures

Several controls are required to assess that our findings are robust to the structure of the RNN, are robust to its input statistics, are robust to other modeling assumptions, and continue to depend on the task being predictive. We describe here a set of controls for the spatial exploration task.

Each control model is trained for a total of 200 epochs, enough for all models to converge. Our focus is not on optimizing performance and therefore we do not employ an Early Stopping Rule here, although we reduce the learning rate on plateau when the validation loss doesn't decrease for more than 10 epochs. We first describe the overall control analysis and detail later the individual models. The key difference for the models is given in their respective names, where we use the abbreviation 'w' for with and 'wo' for without. For example 'wo distance information' refers to the same predictive model trained without distances from the walls in its observations. The models are sorted into three categories: predictive models, non-predictive models and predictive models with critical modifications. These last ones include modifications to the network that differ from the architecture of the main model presented. Some of these have minor differences from the original framework (e.g. adding a sparsity constraint) while others are critically different, like in the case of predicting the previous step (the past instead of the future).

We show how for these models the metrics introduced in the main manuscript - predictive error, latent signal transfer and dimensionality analysis (DG) - capture differences across these models. In Fig. S7 we show how the models converge in their cost function through learning, Fig. S7a. We then show the predictive error symmetry, Fig. S7b based on the position of the axis of symmetry for the predictive error. For a network trained to predict the next step, this should be roughly 1 while for a network trained to predict the previous step, this should be roughly -1. In Fig. S7c we show the linear regression coefficient for a linear regression between the hidden representations of these networks and the spatial variables x, y. The linear decoding of position is the average of the two regressors for coordinate x and y. Models linearly encode for position in their hidden representation have a linear decoding measure closer to one. The results of these controls are in line with those in Fig. 3 of the main manuscript.

Following the same analysis presented in the main manuscript (Fir. 3) we next analyse latent signal transfer, performing a Canonical Correlation Analysis between the position variables and the top 3 Principal Components of the hidden representation for every epoch, Fig. S8a. The same analysis is repeated for the observation signal in Fig. S8b. In predictive models, while the former grows through learning, the latter declines – indicating that the top Principal Components in the hidden representation represent the position (latent signal) rather than observations.

Finally we analyze dimensionality trends across learning for both linear and nonlinear dimensionality measures. Fig. S9a shows the linear dimensionality (PR) across learning while Fig. S9b shows the average of nonlinear dimensionality measures. The trends of predictive and nonpredictive models are highlighted with brackets and generally agree with the trends pointed out in the main manuscript. It is clear that the variability across models is high: these metrics can be affected by several different factors. For example, enforcing sparsity - which can be achieved in several different ways - may modify the dimensionality of the representations. Finally in Fig. S9c we show the dimensionality gain, being the ratio between linear (Fig. S9a) and nonlinear (Fig. S9b dimensionalities.

Having analyzed the metrics introduced in the main manuscript in Figs. S7 to S9 we then turn to the question of place cell coding. As highlighted in the main manuscript the emergence of place cell activation is a possible way to explain and interpret the trends in the metrics established this far. In Fig. S10 we show the place selectivity in the neural activities of 100 neurons across all models. The 100 neurons are sorted to be the 100 neurons with maximum average activity. From this figure it appears that all predictive models develop some form of localized activations while non-predictive models do not. We also aimed at capturing the overall statistics across all cells for their sparsity. We analyzed two forms of sparsity: temporal sparsity and spatial sparsity. For temporal sparsity (Fig. S11a) we compute the average across time of the total activation (L1 norm of activity population vector, given positive activity). For spatial sparsity we compute the average activations of neurons, once such activations have been averaged over space (Fig. S11b). This is the average, for each neuron, of the values shown in Fig. S11a. We also show in Fig. S12 four examples of how different hidden representations appear in PC space. Here the top three Principal Components of the hidden representation are colored by the x-position of the agent in the environment, similar to Fig. 5 in the main text.

Overall the results here displayed confirm the principal results presented in the main manuscript. They also introduce several nuances and avenues for interesting future study. We provide the code to generate these models and analyses.

We now explain the details of the 14 models we compared above. Each model lists only the differences from the original one, which we refer to as "predictive learning."

**Predictive networks**

- Noise in RNN activations. In this model gaussian noise with std of 0.1 is added on top of the activations of every unit at each step.

- Predictive learning with input noise. We add noise to the input as a control that the phenomena describe are not dependent on the absence of noise or on overfitting. We add time independent zero mean gaussian noise to each input channel with an amplitude $\sigma^2_{noise}$ which is 10% of the total variance of the channel: $\sigma_{noise} = 0.1\sigma_{channel}$ for all channels. The model shows the same signatures of predictive learning.

- GRU. In this model the network, instead of being a recurrent vanilla network has units which are GRU.

- LSTM. In this model the network, instead of being a recurrent vanilla network has units which are LSTM.

**Non-predictive networks**

- Autoencoder without bottleneck. In place of the RNN we use a feedforward layer of size 200 and we train the model not on predicting the upcoming observations but rather on replicating them (autoencoding framework). This model can be trained, but it doesn't display all the phenomena highlighted in the main text. The linear dimensionality increases and the intrinsic dimensionality decreases but the latent variables do not seem to be extracted as in the predictive case. Both CCA metrics fail to show the extraction of latent variables and place cell tuning curves do not appear.

- Autoencoder with bottleneck. In place of the RNN we use a feedforward layer of size 10.

- Non-predictive, recurrent autoencoder. This model, as discussed in the main text, has the same structure of the predictive learning one but is trained in replicating (autoencoding) the input observations.

#### Other models

- Sparsity, predictive learning with a sparsity constraint. We add a L1 sparsity constraint with a penalty of $5\ 10^{-8}$ on the activations of the recurrent network. This constraint doesn't appear to sparsify the network in a straightfoward way. Rather it seems to strongly reduce the overall activity and introduce a code where some units tend to be more active than others overall. This is a signature of a less distributed neural code.

- Predictive learning without actions. In this case actions are not fed as input to the network but the network is still trained to reproduce both distance and color information. The task is more difficult but the network still seems to be able to extract a representation with similar features to the full case, as long as it is trained to perform predictive learning.

- Predictive learning without color information. We train the same predictive learning model without color information from the sensors in the input and output: color information is not passed as input and is not decoded from the output. Sensors receive only distance information. The model minimizes the cost function but it doesn't display the features analyzed in the main text.

- Predictive learning without distance information: same as above but without distance information. The model seems to learn with similar characteristics, showing robustness to the lack of distance information. This is an important feature as one may say that having precise, "hard-coded" distance information in the sensors is not biological. In the main text we study the case of both distance and color information to include reasonably available visual information, but the present control is important to highlight the robustness of our results.

- Autoencoder with angle. We train a network to autoencode its observations where to the observations the current angle of the agent is added. The model didn't train particularly well across several repetitions we tried; it is included as an example of model which fails to train in outputting the angle, as compared to other autoencoding models listed above.

- Predictive learning on the previous timestep. We train the same model but to predict the previous time step in time rather than the future one. This model doesn't extract the latent space.

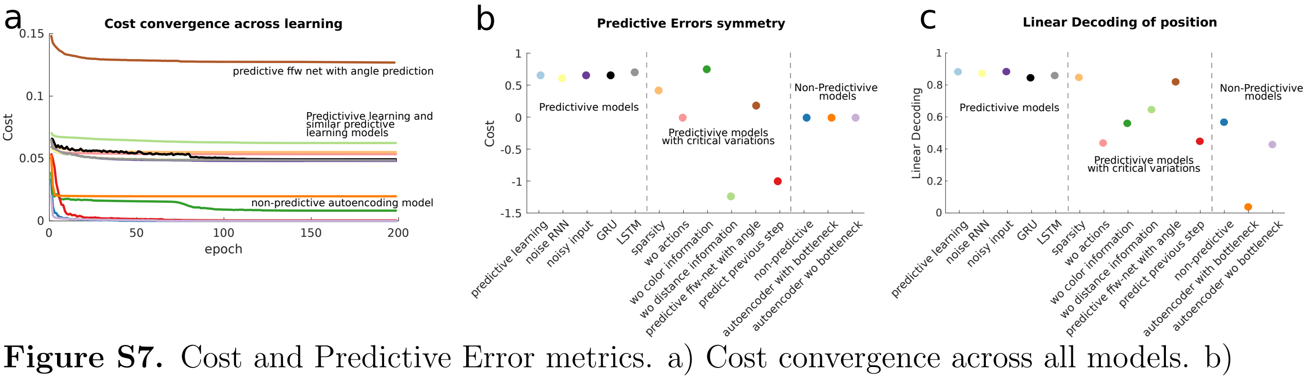

**Figure S7.** Cost and Predictive Error metrics. a) Cost convergence across all models. b) Predictive error symmetry axis position upon learning across all models. This is the same measure used for Fig. 4 main manuscript. c) Linear Decoding performance of the latent variables, from the network representation.

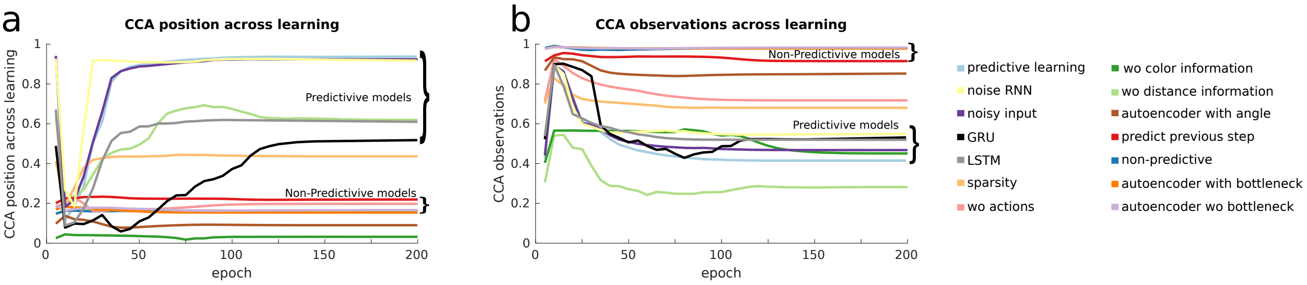

**Figure S8.** Canonical Correlation Analysis. a) Canonical correlation analisys through learning between the top 3 PCs of the hidden representation and the position of the agent in (x,y) coordinates. The average of the two canonical correlations found by the analysis is plotted for each epoch. b) Same as panel a but between the hidden representation and the top 3 PCs of the observations.

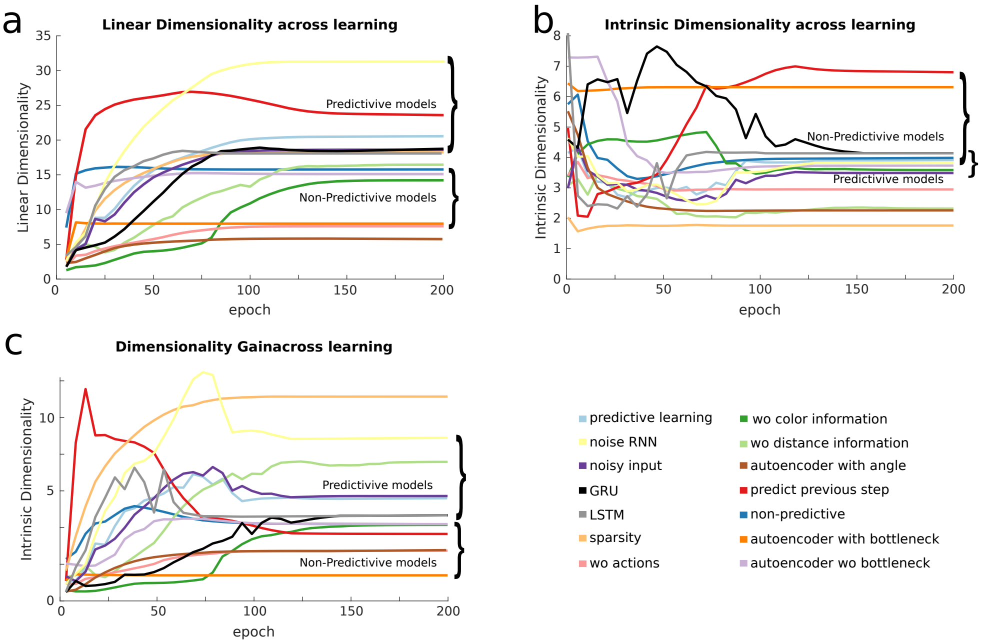

**Figure S9.** Dimensionality analysis. a) Linear dimensionality (PR) across learning. b) Non-linear dimensionality (ID) across learning. Each curve is the average of the 4 ID estimation methods introduced in the main manuscript, cf. Fig. 3. c) Dimensionality Gain across learning.

### 4 Pilot analysis of neural data

#### 4.1 Neural data reveal partial evidence of predictive learning.

We analyze a publicly available neural dataset [14], collected in the Buszaki lab, consisting of recordings from the hippocampal area CA1. In the analyzed session I15, rat i01 performed free exploration of an open square environment for about 60min. Over 102 channels 164 CA1 neurons were recorded and identified. We didn't preprocess the data except for binning spikes into a moving window of 100ms (we repeated the procedure for a moving window of 50ms and obtained similar results).

First, we decoded the future and past position and head direction of the animal from the neural population at the current time, as a function of the time difference Delta $t$. Fig. S13a shows that the decoding of the spatial coordinates, but not the angle, appeared to be prospective in time by about 100ms. This result is in line with our findings regarding predictive error and decoding of latent variables from the representation, cf. Figs. 3a,b.

We then fit, by means of a quadratic Generalized Linear Model, receptive fields to each neuron. We repeated the same procedure both in the spatial domain of the environmental coordinates

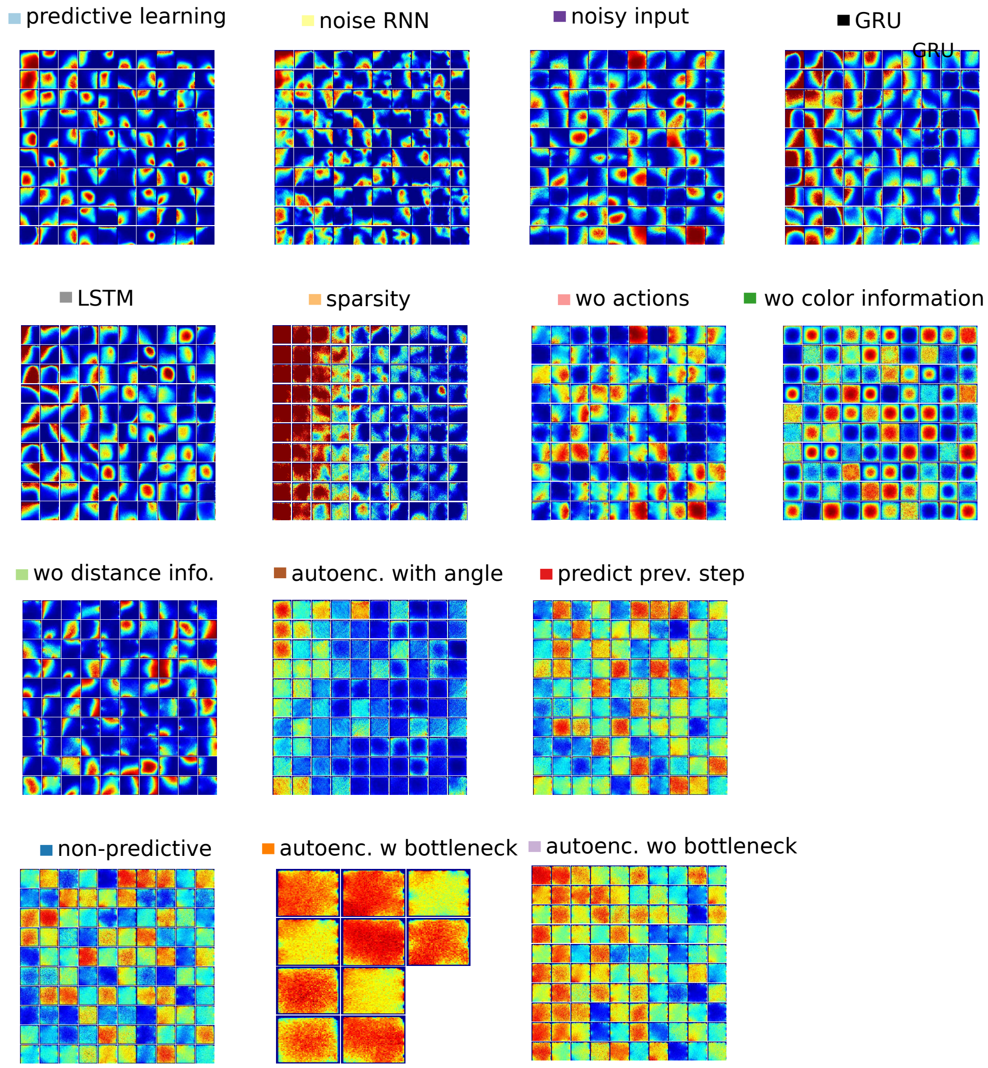

**Figure S10.** Average activity of 100 neurons in the latent (environment) space across all models.

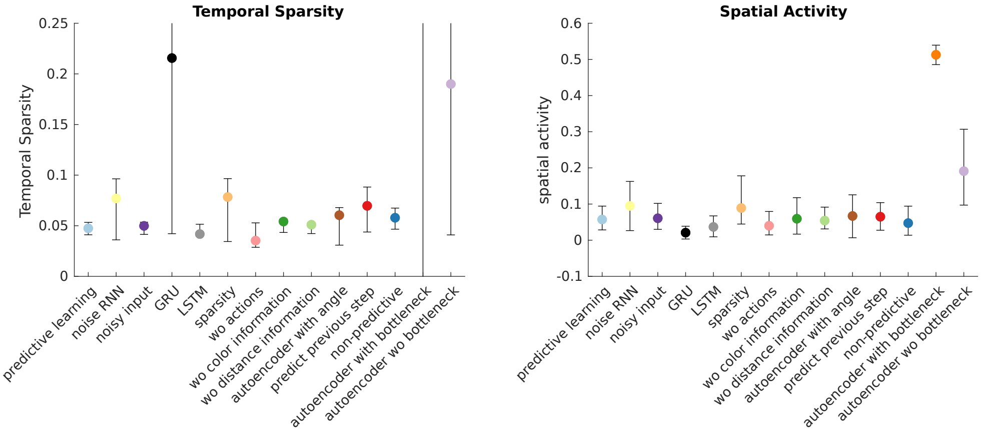

**Figure S11.** Sparsity analysis. a) Temporal sparsity. Average L1 norm of the population vector across time. For each time step the L1 norm of the activity of all neurons is computed and the mean and standard deviation of the distribution of such sparsity measure are displayed. b) Spatial sparsity. We compute the L1 norm of the spatial averages of individual neurons. These are the ones plotted in Fig. S10. The average and standard deviation of the distributions of L1 norms are used for the plot across all models.

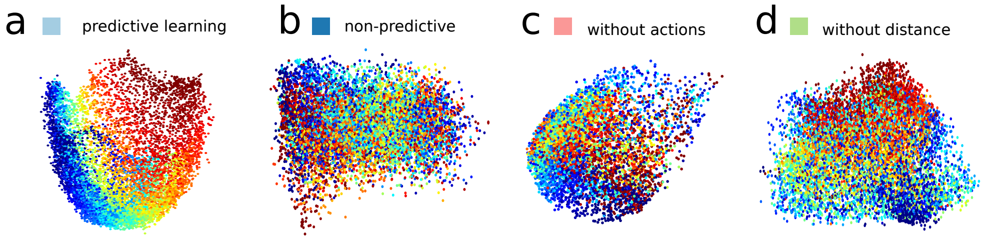

**Figure S12.** Principal Component Space. a) We show the hidden representation in PCs1-2 colored by the x coordinate of the latent space (environment). This is the same plot as in Fig. 5 of the main manuscript. b) Same as panel a for the non-predictive model. c) Same as panel a for the predictive model without actions. d) Same as panel a for the predictive model without distance information.

and in the Principal Component space spanned by the first two PCs. We measured the size of the field by the negative exponent of the fit constant (exponential decay of the field). A higher exponent indicates a faster decay and thus a sparser code, Fig. S13b. Neural receptive fields fit to spatial (blue) and PCs coordinates (red) have a generally similar form. This result is in line with our analysis developed in the main manuscript Fig. 4. There we showed how, in our simulations, single neurons developed localized receptive fields on the neural population (PC) manifold.

Finally, we tested different measures of intrinsic dimensionality on the neural data, in Fig. S13c. We reported only measures which displayed numerical stability. Interestingly, several measures which appeared very stable in simulations seemed unstable on neural data. This could be due to the lack of data (e.g. the green curve in Fig. S13c appears numerically stable when at least 40 neurons are used for the computation), or to the intrinsic noise of neural data (which was not modeled in our simulations). This suggests that careful future analyses are needed to understand the problem of estimating the dimensionality of neural data, a topic which recent work suggests could be of crucial importance to understand hippocampal coding [10].

We close by pointing out that the metrics we developed (latent signal transfer and dimensionality analysis) are mainly geared towards understanding learning and the formation of manifold structures through the learning process (Fig. 3 main manuscript). We thus look forward to future analysis of datasets with more neurons and to attendant tests of how our methods may reveal how the geometrical properties of neural representations evolve through task learning.

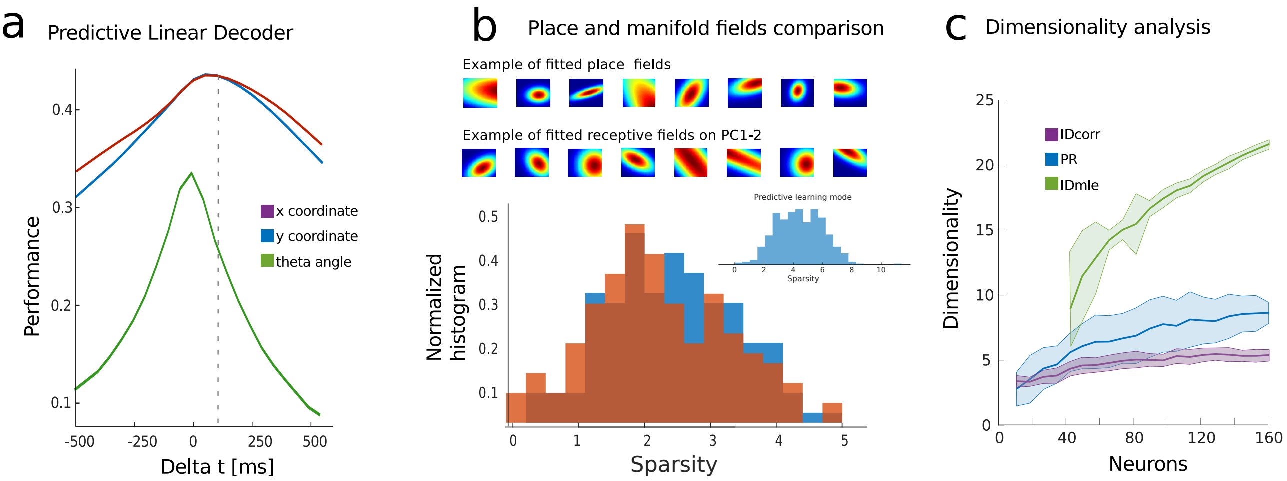

**Figure S13.** Neural data analysis. a) Linear decoding of latent variables from neural population data. b) Comparison between localization of receptive fields in the latent space vs Principal Components space. Top example of localization of extracted receptive fields (quadratic GLM model) on both latent space variables (x,y) and PCs 1-2. Bottom comparison of the extracted tuning in the two cases (red for fitted place fields and blue for fitted fields on PCs). Inset: distribution for the predictive learning model. c) Three measures of dimensionality estimation applied to neural data.

5. F. Camastra and A. Staiano, *Intrinsic dimension estimation: Advances and open problems*, Information Sciences, 328 (2016), pp. 26–41.

6. H. Dai, Z. Geary, and L. P. Kadanoff, *Asymptotics of eigenvalues and eigenvectors of toeplitz matrices*, Journal of Statistical Mechanics: Theory and Experiment, 2009 (2009), p. P05012.

7. P. Gao, E. Trautmann, B. M. Yu, G. Santhanam, S. Ryu, K. Shenoy, and S. Ganguli, *A theory of multineuronal dimensionality, dynamics and measurement*, bioRxiv, (2017), p. 214262.

8. S. Lahiri, P. Gao, and S. Ganguli, *Random projections of random manifolds*, arXiv preprint arXiv:1607.04331, (2016).

9. A. Litwin-Kumar, K. D. Harris, R. Axel, H. Sompolinsky, and L. F. Abbott, *Optimal Degrees of Synaptic Connectivity*, Neuron, 93 (2017), pp. 1153–1164.e7.

10. R. J. Low, S. Lewallen, D. Aronov, R. Nevers, and D. W. Tank, *Probing variability in a cognitive map using manifold inference from neural dynamics*, bioRxiv, (2018).

11. J. Mairal, F. Bach, J. Ponce, and G. Sapiro, *Online dictionary learning for sparse coding*, in Proceedings of the 26th annual international conference on machine learning, ACM, 2009, pp. 689–696.

12. L. Mazzucato, A. Fontanini, and G. La Camera, *Stimuli Reduce the Dimensionality of Cortical Activity*, Frontiers in Systems Neuroscience, 10 (2016).

13. J. O'Keefe and J. Dostrovsky, *The hippocampus as a spatial map. Preliminary evidence from unit activity in the freely-moving rat.*, Brain Res, 34 (1971), pp. 171–175.

14. E. Pastalkova, Y. Wang, K. Mizuseki, and G. Buzsáki, *Simultaneous extracellular recordings from left and right hippocampal areas ca1 and right entorhinal cortex from a rat performing a left/right alternation task and other behaviors*, CRCNS, (2015).

15. C. Pehlevan, A. M. Sengupta, and D. B. Chklovskii, *Why do similarity matching objectives lead to hebbian/anti-hebbian networks?*, Neural computation, 30 (2018), pp. 84–124.

16. M. Rezghi and L. Elden, *Diagonalization of tensors with circulant structure*, Linear Algebra and its Applications, 435 (2011), pp. 422–447.

17. A. Sengupta, M. Tepper, C. Pehlevan, A. Genkin, and D. Chklovskii, *Manifold-tiling localized receptive fields are optimal in similarity-preserving neural networks*, bioRxiv, (2018).

18. T. Solstad, C. N. Boccara, E. Kropff, M.-B. Moser, and E. I. Moser, *Representation of Geometric Borders in the Entorhinal Cortex*, Science, 322 (2008), pp. 1865–1868. WOS:000261799400061.

19. H. Stensola, T. Stensola, T. Solstad, K. Froland, M.-B. Moser, and E. I. Moser, *The entorhinal grid map is discretized*, Nature, 492 (2012), pp. 72–78. WOS:000311893400047.

20. T. J. Wills, F. Cacucci, N. Burgess, and J. O'Keefe, *Development of the Hippocampal Cognitive Map in Preweanling Rats*, Science, 328 (2010), pp. 1573–1576. WOS:000278859200051.
